# IGFBP2 expressing midlobular hepatocytes preferentially contribute to liver homeostasis and regeneration

**DOI:** 10.1101/2022.10.12.511675

**Authors:** Yu-Hsuan Lin, Yonglong Wei, Yunguan Wang, Chase A. Pagani, Lin Li, Min Zhu, Zixi Wang, Meng-Hsiung Hsieh, Yu Zhang, Tripti Sharma, Tao Wang, Hao Zhu

**Affiliations:** Children’s Research Institute, Departments of Pediatrics and Internal Medicine, Center for Regenerative Science and Medicine, University of Texas Southwestern Medical Center, Dallas, TX 75390, USA; Quantitative Biomedical Research Center, Department of Population and Data Sciences, Center for the Genetics of Host Defense, University of Texas Southwestern Medical Center, Dallas, TX, 75390, USA

**Keywords:** Zone 2, Lineage tracing, Zonation, Midlobular hepatocytes, Regeneration

## Abstract

While midlobular hepatocytes in zone 2 are a recently identified cellular source for liver homeostasis and regeneration, these cells have not been exclusively fate mapped. We generated a *Igfbp2-CreER* knockin strain, which specifically labels midlobular hepatocytes. During homeostasis over 1 year, zone 2 hepatocytes increased in abundance from occupying 21% to 41% of the lobular area. After either pericentral injury with carbon tetrachloride or periportal injury with DDC, IGFBP2+ cells replenished lost hepatocytes in zones 3 and 1, respectively. IGFBP2+ cells also preferentially contributed to regeneration after 70% partial hepatectomy, as well as liver growth during pregnancy. Because IGFBP2 labeling increased substantially with fasting, we used single nuclear transcriptomics to explore zonation as a function of nutrition, revealing that the zonal division of labor shifts dramatically with fasting. These studies demonstrate the contribution of IGFBP2-labeled zone 2 hepatocytes to liver homeostasis and regeneration.

**Highlights:**

- Lineage tracing showed that midlobular hepatocytes proliferate during homeostasis.
- Zone 2 cells were protected from portal and centrilobular injuries and replaced lost hepatocytes.
- Fasting induced significant changes in liver zonation.

**eTOC statement:** The liver consists of different zones with spatial heterogeneity in their metabolic functions. Here, a new *Igfbp2-CreER* line enabled direct tracing of midlobular hepatocytes, which showed that IGFBP2+ cells serve as a source of new hepatocytes during normal homeostasis and regeneration.

## Introduction

The source of new liver cells during homeostasis and regeneration has been intensely debated. Many studies have proposed diverse cell types as liver stem cells: Sox9 expressing cells near the portal triad (Font-Burgada et al., 2015; Furuyama et al., 2011); Axin2 expressing hepatocytes near the central vein (Wang et al., 2015); Tert expressing hepatocytes distributed throughout the lobule (Lin et al., 2018). Other studies have challenged these claims (Sun et al., 2020)(Chen et al., 2020). As single cell technologies were applied to the liver, metabolic zonation emerged as an important source of cellular heterogeneity (Ben-Moshe and Itzkovitz, 2019). To determine whether this zonal heterogeneity leads to differences in regenerative capacity, we previously compared the contraction and expansion of different zonal compartments using 14 orthogonal *CreER* mouse strains. Using these tools, we and others showed that zone 2 cells are a major source of new hepatocytes during liver growth, maintenance, and tissue repair (Chen et al., 2020; He et al., 2021; Wei et al., 2021).

While midlobular hepatocytes in zone 2 are a potential source of new hepatocytes, there are still major questions and unresolved controversies. Prior studies, including ours, used negative tracing approaches to generate support for zone 2 expansion. Because zone 2 cells are ill-defined and do not have a singular genetic marker, these studies were unable to exclusively trace zone 2 cells. For example, we used *Hamp2*, which is enriched in zone 2, to probe this population. However, *Hamp2* is also expressed in some zone 1 and 3 cells. Other studies using pan-zonal AAV-TBG-Cre and Ki-67-CreER tracing tools faced similar caveats (Chen et al. 2020; He et al. 2021). In the current study, we focused on a maker that is more restricted to zone 2 cells. We generated a *Igfbp2-CreER* knockin strain that specifically labels midlobular hepatocytes. Our study of the *Igfbp2*+ population again showed that zone 2 populations preferentially contribute to homeostatic repopulation of the liver over long time periods. In addition, our studies comparing Igfbp2 to other markers such as Hamp2 suggest heterogeneity among different zone 2 populations.

Because there were previously no exclusive tracing markers for zone 2, there was no high fidelity method to assess zone 2’s contribution to diverse regenerative processes. Here, we used the *Igfbp2* tracer to independently assess the contribution of zone 2 cells in both commonly and poorly studied injury assays. Our results provided direct evidence that midlobular cells regenerate after zone 1 and 3 injuries caused by liver toxins. In addition, we quantified how much zone 2 cells contribute to 1) partial hepatectomy induced regeneration, 2) hepatocyte transplant induced repopulation of a chronic liver disease model, and 3) a profound but rarely studied process - pregnancy induced liver enlargement. Remarkably, the preferential contribution from zone 2 was observed in all of these models.

## Results

### *Igfbp2-CreER* mice efficiently label midlobular hepatocytes

Using CRISPR genome editing, we created a new *Igfbp2-CreER* strain in which the IRES cassette followed by *CreERT2* recombinase was integrated into the endogenous 3’ untranslated region (3’UTRs) of *Igfbp2* (**Figure 1A**). By crossing *Igfbp2-CreER* and *Rosa-LSL-tdTomato* reporter mice, we generated lineage tracing mice with permanent Tomato protein expression in IGFBP2+ hepatocytes after tamoxifen administration (**Figure 1B**,**C** and **S1A**,**B**). There was very little reporter activation (less than 0.1%) when oil as a vehicle was given, and this rare population did not significantly contribute to homeostasis after 26-weeks of tracing (**Figure S1C**). We quantified the distribution of Tomato+ cells across all zones and found that it was highest in the midlobule, or zone 2 (**Figure 1D**). Almost no Glutamine Synthetase (GS)+ cells in zone 3 of the lobule were labeled by Tomato. Occasional cells adjacent to the portal trials were labeled, but the labeling in this zone 1 region was much more sparse than in zone 2.

**Figure 1.**
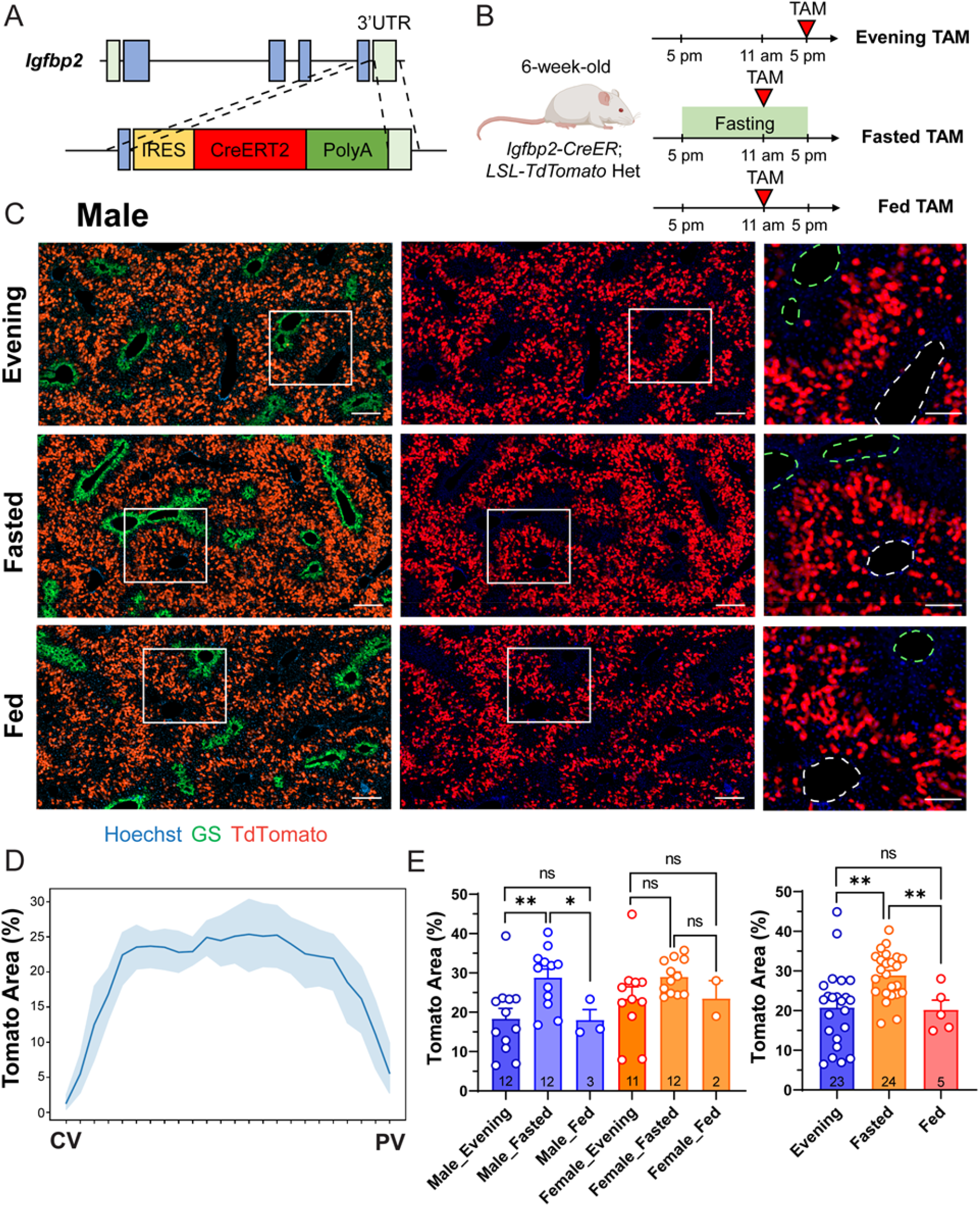
The *Igfbp2-CreER* strain labels zone 2 hepatocytes in a nutrient dependent manner. **A**. Schema of the homology directed repair (HDR) construct used to generate the *Igfbp2-CreER* strain. **B**. Schema of three different tamoxifen administration approaches. **C**. Representative images of male *Igfbp2-CreER; tdTomato* het mouse livers two weeks after tamoxifen given at 100mg/kg intraperitoneally using different approaches. In magnified images, the green dashed circles represent CVs (marked by GS), and the white dashed circles represent PVs. Scale bar = 200 μm for cropped images and 100 μm for magnified images. **D**. The percent area labeled in *Igfbp2-CreER* mice given evening tamoxifen. **E**. The quantification of the Tomato+ areas of the three tamoxifen approaches. The right panel combines the data points from the two sexes shown in the left panel. The data points of evening and fasting tamoxifen are the same as in the 2-week timepoint in **Figure 2C** and **S4D** (n = 23, 24 and 5 mice for evening, fasted, and fed groups respectively).

*Igfbp2* expression is known to be nutritionally regulated, as hepatic *Igfbp2* mRNA increases after fasting (Tseng et al., 1992). To examine whether Cre mediated labeling efficiency was also nutritionally regulated, mice were given tamoxifen at noon after overnight fasting (**Figure 1B-C**). To exclude the potential impact from circadian regulation, control mice that were fed normally were also given tamoxifen at noon. Compared to fed mice given tamoxifen in the evening or fed mice given tamoxifen at noon (20.79 ± 9.75% and 20.2 ± 5.47% (mean ± SD) respectively), overnight fasted mice had a large increase in Tomato labeling (28.90 ± 5.93%) (**Figure 1E**). The fasted group also had reduced variability in Tomato labeling than the fed group given tamoxifen in the evening.

Genomic *CreER* integration can cause unexpected biological effects that can influence hepatocyte turnover and long term lineage tracing results. To confirm that the *Igfbp2-CreER* transgene did not influence hepatocyte turnover, we gave the mice 5-ethynyl-2′-deoxyuridine (EdU) in the drinking water for 10 days (**Figure S2A**). The frequency and spatial distribution of EdU+ hepatocytes were quantified as previously described (Lin et al., 2018). Compared to wild-type (WT) livers, *Igfbp2-CreER* knockin mice exhibited a similar EdU distribution with zone 2 having the highest frequency of proliferating cells (**Figure S2B**). Collectively, these results showed that the *Igfbp2-CreER* strain can efficiently label midlobular hepatocytes to different extents with the evening or fasting tamoxifen approach. Furthermore, there was no evidence that this strain caused significant ectopic labeling or artificially flavored clonal expansion from any particular zone.

### Midlobular zone 2 cells marked by *Igfbp2* expand during normal homeostasis

We previously used distinct *CreER* lines to label different zonal compartments and systematically compared the contraction and expansion between different hepatocytes subpopulations (Wei et al., 2021). The results revealed that zone 2 hepatocytes are the major source for renewal during normal homeostasis. Sparsely labeled zone 2 hepatocytes by *Hamp2-CreER* (7.4 ± 0.7% of the slide area was labeled) showed clonal expansion after 12 months tracing (27.4 ± 4.1%) (Wei et al. 2021). To determine if the expansion was only applicable to the rare subpopulation labeled by HAMP2, we employed the *Igfbp2-CreER* line for lineage tracing under steady-state homeostatic conditions. Fed mice were given tamoxifen in the evening and traced for 2, 12, 26 and 52 weeks (**Figure 2A, S3A**). The images from all timepoints exhibited a similar zonated pattern of Tomato distribution, indicating that there was no replacement of unlabeled populations from either zone 1 or zone 3 (**Figure 2B, S3B**). To evaluate whether labeled zone 2 cells increased in number during homeostasis, we measured the area occupied by Tomato+ cells over 12-, 26- and 52-week periods. In the evening tamoxifen group, the Tomato labeled area went from 20.79 ± 9.75% (2 weeks post-tamoxifen) to 31.13 ± 14.47% (12 weeks), 34.79 ± 14.63% (26 weeks), and 40.71 ± 14.96% (52 weeks) (**Figure 2C**).

**Figure 2.**
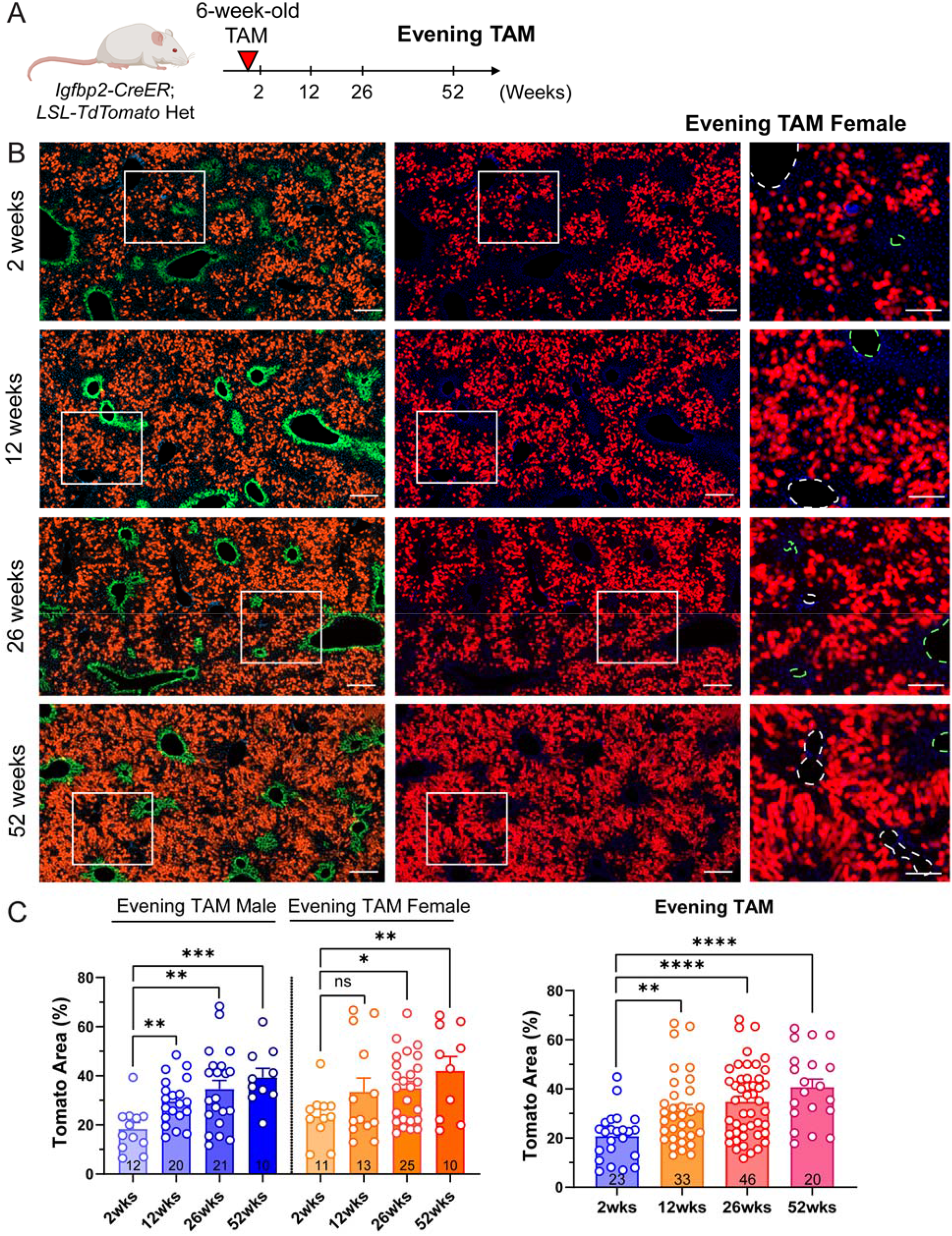
Zone 2 hepatocytes give rise to new hepatocytes during normal homeostasis. **A**. Schema of evening tamoxifen lineage tracing experiment under normal homeostasis. **B**. Representative images of female evening tamoxifen lineage tracing over 2, 12, 26 and 52 weeks under homeostatic conditions. Scale bar, 200 μm for cropped images and 100 μm for magnified images. Slides were stained for GS (green). The green dashed circles represent CVs (marked by GS), and the white dashed circles represent PVs. **C**. Quantification of the Tomato area from **B**. The right panel combines the data points from two sexes shown in the left panel. The 2-week data points are the same as the evening timepoints in **Figure 1E** (n = 23, 33, 46 and 20 mice for 2, 12, 26 and 52 weeks).

Because zone 2 has been defined differently in various studies, the extent to which midlobular populations expand is dependent on the size of the zone 2 population labeled initially. We reasoned that increasing the initially labeled size of the zone 2 domain would lead to a reduction in the relative expansion over time. To label and trace a broader midlobular population, we performed the homeostasis tracing experiments using tamoxifen labeling under fasting conditions. Similar spatial distributions of Tomato+ cells were observed over 2, 12 and 26 weeks (**Figure S4A-C**). We noticed an increase of Tomato area from 28.90 ± 5.93% (2 weeks) to 35.26 ± 12.32% (12 weeks) and 37.27 ± 10.05% (26 weeks) after fasting tamoxifen labeling (**Figure S4D**). Compared to evening tamoxifen, the change in the Tomato+ area was lower in the fasting labeled group. This suggested that some of the additionally labeled cells near the PV likely did not contribute to the expansion, since the final tomato labeling result was similar. Overall, we found that midlobular zone 2 hepatocytes labeled by *Igfbp2-CreER* line expanded during homeostasis.

### GFBP2 expressing zone 2 cells contribute to regeneration after zone 1 and 3 injuries

To determine whether zone 2 cells labeled using *Igfbp2-CreER* mice contribute to regeneration after common liver injuries, we challenged mice with toxins that damage hepatocytes in zone 1 and 3 (**Figure 3A, 3D**). In these experiments, we used the fasting tamoxifen approach to increase the consistency of labeling before various injuries. After tamoxifen labeling, we injected mice with either single doses of carbon tetrachloride (CCl_4_) or acetaminophen (APAP) to cause centrilobular hepatic necrosis. After two weeks, the necrotic areas fully recovered and were replaced by the remaining adjacent hepatocytes. Tomato+ cells shifted from the midlobular to centrilobular regions, even coexpressing Tomato and GS, indicating that proliferating zone 2 hepatocytes regenerated to replace the cells lost in zone 3 (**Figure 3B**). The percentage of Tomato labeling was 50.24 ± 9.95% and 49.52 ± 10.57% after one dose CCl_4_ or APAP, compared to non-injured control mice (32.75 ± 6.16%) (**Figure 3C**). To investigate a cholestatic injury focused on the opposite end of the lobule, we fed mice with 0.1% 3,5-diethoxycarbonyl-1,4-dihydrocollidine (DDC) for 4 weeks. Chronic DDC feeding increases biliary porphyrin secretion and induces a ductular reaction, modeling human cholangiopathies such as primary sclerosing cholangitis, primary biliary cirrhosis, and drug-induced bile duct damage (Fickert et al., 2007). After DDC, biliary damage was associated with ductular reactions in periportal regions, which were surrounded by Tomato+ cells (**Figure 3E**). To allow the recovery and regeneration of hepatocytes, we discontinued DDC for two weeks. After this washout period, the periportal area was filled with Tomato+ cells (**Figure 3E**). The Tomato+ area was 39.44 ± 6.07% in DDC fed mice, compared to non-injured control mice (33.45 ± 3.73%) and to injured mice before washout (31.58 ± 2.62%) (**Figure 3F**). Since a majority of liver injuries cause damage in zone 1 or 3, zone 2 hepatocytes are well positioned to avoid destruction. Altogether, these data suggested that the midlobular cells are capable of expansion, and ultimately regenerate the liver after common liver injuries.

**Figure 3.**
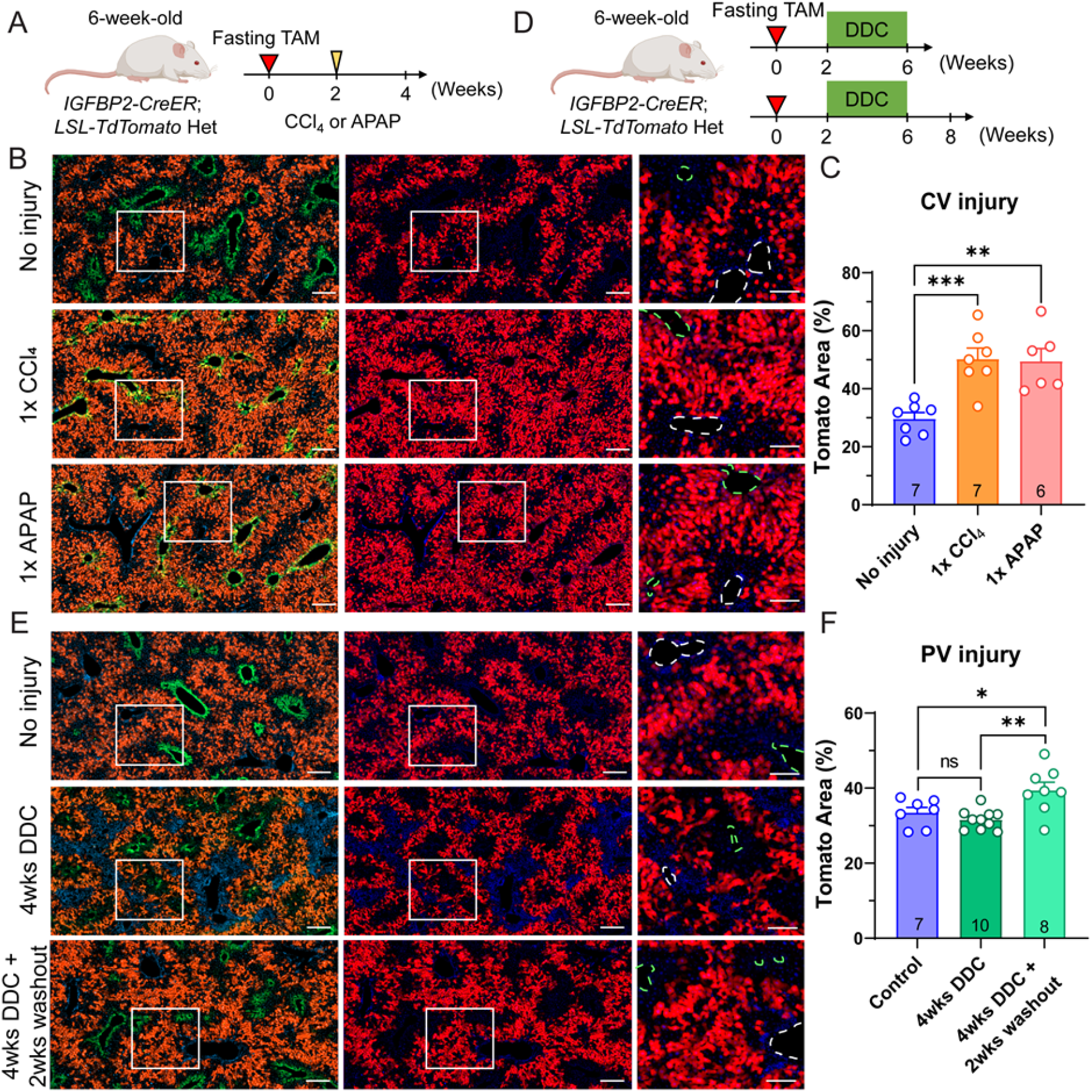
*Igfbp2*+ zone 2 cells regenerate after both pericentral and periportal injuries. **A**. Schema of fasting tamoxifen male challenged with centrilobular chemical injury CCl_4_ or APAP. **B**. Representative images of fasting tamoxifen treated males without injury or two weeks after CCl_4_ or APAP injury. Scale bar, 200 μm for cropped images and 100 μm for magnified images. Slides were stained for GS (green). The green dashed circles represent CVs (marked by GS), and the white dashed circles represent PVs. **C**. Quantification of the Tomato area from **B** (n = 7, 7 and 6 mice for no injury, CCl_4_ and APAP, respectively). **D**. Schema of fasting tamoxifen male given 4 weeks of DDC diet. **E**. Representative images of fasting tamoxifen male without injury, after 4 weeks of DDC feeding, or after 4 weeks DDC plus a 2 week washout period with normal chow. Scale bar, 200 μm for cropped images and 100 μm for magnified images. Slides were stained for GS (green). The green dashed circles represent CVs (marked by GS), and white dashed circles represent PVs. **F**. Quantification of the Tomato area from **E** (n = 7, 10 and 8 mice for control diet, 4 weeks of DDC, and 4 weeks of DDC and 2 weeks of washout).

### IGFBP2+ zone 2 cells contribute to regeneration after partial hepatectomy

To assess the hepatocyte subpopulations that contribute to liver regrowth, we performed partial hepatectomy (PHx) on our lineage tracing mice. Compared to chemical injuries, PHx is not known to cause zone-specific damage. We used both the evening and fasting tamoxifen approaches to lineage trace after different extents of zone 2 labeling. To examine zone 2 hepatocyte contributions, we compared the Tomato labeling between two liver lobes obtained from the initial surgical resection and the remaining lobes 14 days after surgery (**Figure 4A**). Pre- and post-PHx livers showed a similar spatial distribution of Tomato+ cells (**Figure 4B, S5B**). This comparable zonal distribution revealed that there were contributions to regeneration from all three zones, in contrast to what was observed with pericentral or periportal injuries (**Figure 3B, 3E**). Indeed, it is known that most hepatocytes across the lobule proliferate after surgical resection (He et al., 2021). To control for the variable labeling that occurs between individual mice, we further quantified the change in Tomato labeling frequency within each individual mouse, before and after surgery. This revealed a small but significant increase in Tomato labeling from 21.88 ± 11.25% to 26.27 ± 12.36% (**Figure 4C**). Similar results were also obtained from the fasting tamoxifen approach (27.29 ± 6.51% to 29.11 ± 8.41%) (**Figure S5C-D**). Previous studies showed that zone 1 and zone 2 hepatocytes together contribute more than zone 3 to liver regeneration after PHx (He et al. 2021; Chembazhi et al. 2021), and our results showing a preferential contribution from zone 2 support those observations.

**Figure 4.**
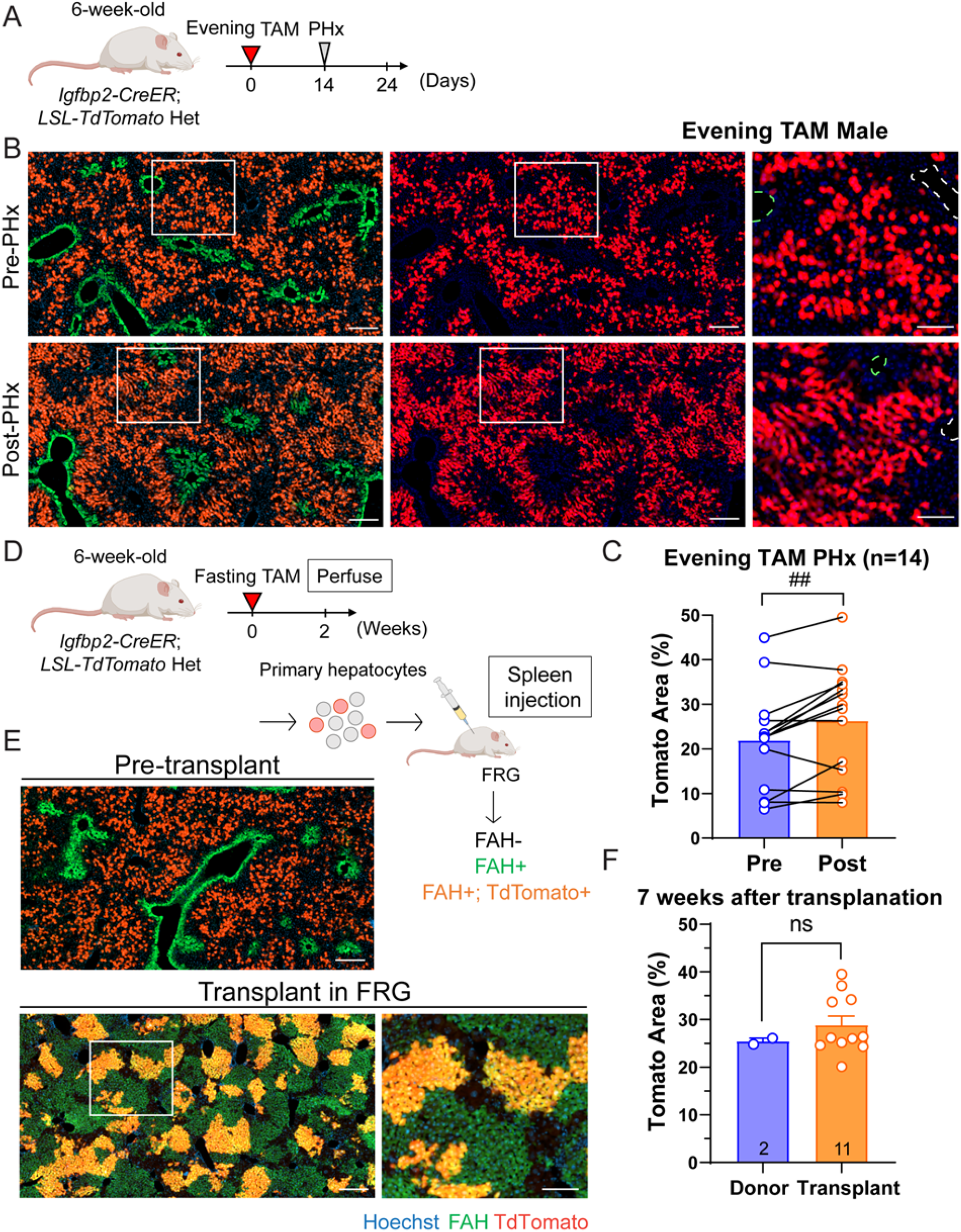
Zone 2 hepatocytes labeled by *Igfbp2-CreER* contribute to liver regeneration and repopulation. **A**. Schema of the *Igfbp2-CreER* lineage tracing experiment in the context of PHx. **B**. Representative images of evening tamoxifen male livers pre- and post-PHx. Scale bar, 200 μm for cropped images and 100 μm for magnified images. Slides were stained for GS (green). The green dashed circles represent CVs (marked by GS), and the white dashed circles represent PVs. **C**. Quantification of the Tomato area from **B**. Significance was assessed by paired t-tests (n = 14 mice). **D**. Schema of primary hepatocyte transplantation from *Igfbp2-CreER* mice into FRG mice. **E**. Representative images of pre-transplant donor livers and the post-transplant repopulated livers. Scale bar, 200 μm for cropped images and 100 μm for magnified images. In the donor livers, slides were stained for GS (green). In the transplanted FRG livers, slides were stained for FAH (green). **F**. Quantification of the Tomato area from **E**. The percentage was quantified by taking the ratio of the Tomato area over the FAH area in the transplant livers (n = 2 and 11 mice for the donors and transplant recipients).

We next asked if hepatocytes from particular zones might serve as better donors in a hepatocyte transplant model. To do this, we transplanted zone 2 hepatocytes into immunodeficient Fah^-/-^ mice (Fah^-/-^/ Rag2^-/-^/ Il2rg^-/-^ [FRG]), a model of tyrosinemia type I that can serve as a transplant recipient (Azuma et al., 2007). We used *Igfbp2-CreER* mice after fasting tamoxifen as the source of donor hepatocytes. To minimize technical bias from flow cytometry, we perfused *Igfbp2-CreER; LSL-tdTomato* livers and without sorting, directly transplanted one million viable hepatocytes into FRG recipients (**Figure 4D**). In this way, a mixture of primary hepatocytes that were Tomato+ (zone 2) and Tomato- (zone 1 and zone 3) populations could be transplanted into FRG recipients through splenic injection. The experiment was done with 7 FRG mice receiving donor #1 hepatocytes and 4 FRG mice receiving donor #2 hepatocytes. The left lateral lobes from each *Igfbp2-CreER* donor were removed before perfusion to ascertain the labeling efficiency of the midlobular population (**Figure 4E**). The livers were harvested 7 weeks after transplantation and were immunostained for FAH to characterize the repopulation by all donor cells (**Figure 4E**). We observed patches of Tomato+ cells clustered together, indicating clonal expansion by donor derived cells. After repopulation, we calculated the percentage of zone 2 cells using the ratio of Tomato+ image area over FAH+ image area. There was a trend toward Tomato percentage increase detected between the donors (25.45 ± 0.92%) and the recipients (28.85 ± 6.19%) (**Figure 4F**). These results suggest that zone 2 versus zone 1 and 3 hepatocytes have a very similar capacity to contribute to liver repopulation after transplantation. If there was a rare cell population dominating repopulation, the fraction of Tomato areas should dramatically increase or decrease. The patches of Tomato+ cells suggest that clonal expansion occurred, but it is unlikely to have been stem cell self-renewal since the Tomato negative cells also expanded. Both the PHx and FRG transplantation models demonstrated a high regenerative capacity of most hepatocytes that can promptly proliferate after loss of hepatic tissue.

### Zone 2 hepatocytes contribute to pregnancy induced liver expansion

To meet the increased metabolic demands of pregnancy, the maternal liver grows and dramatically increases in size (Dai et al., 2011). Because this phenomenon is not frequently studied, it is unknown if there are differences in zonal contributions to pregnancy induced liver expansion. We utilized the *Igfbp2-CreER* line with the fasting tamoxifen approach to trace zone 2 hepatocytes during pregnancy. As a washout period, we waited four weeks after tamoxifen administration before mating was initiated. After delivery, offspring were separated from the mothers on the day of birth (P0) (**Figure 5A**). Remarkably, the body weight, liver weight, and liver to body weight ratios were all significantly increased in the pregnant cohort (**Figure 5B-C**). These data indicate that liver growth increases much more than would be expected if the liver was just enlarging to keep up with body weight changes. The image and quantification of non-mated control females (34.37 ± 7.88%) and P0 mothers (41.81 ± 7.82%) showed an increase in Tomato labeling (**Figure 5D, 5E**), indicating that zone 2 hepatocytes are preferentially contributing to pregnancy induced liver growth more than other zones.

**Figure 5.**
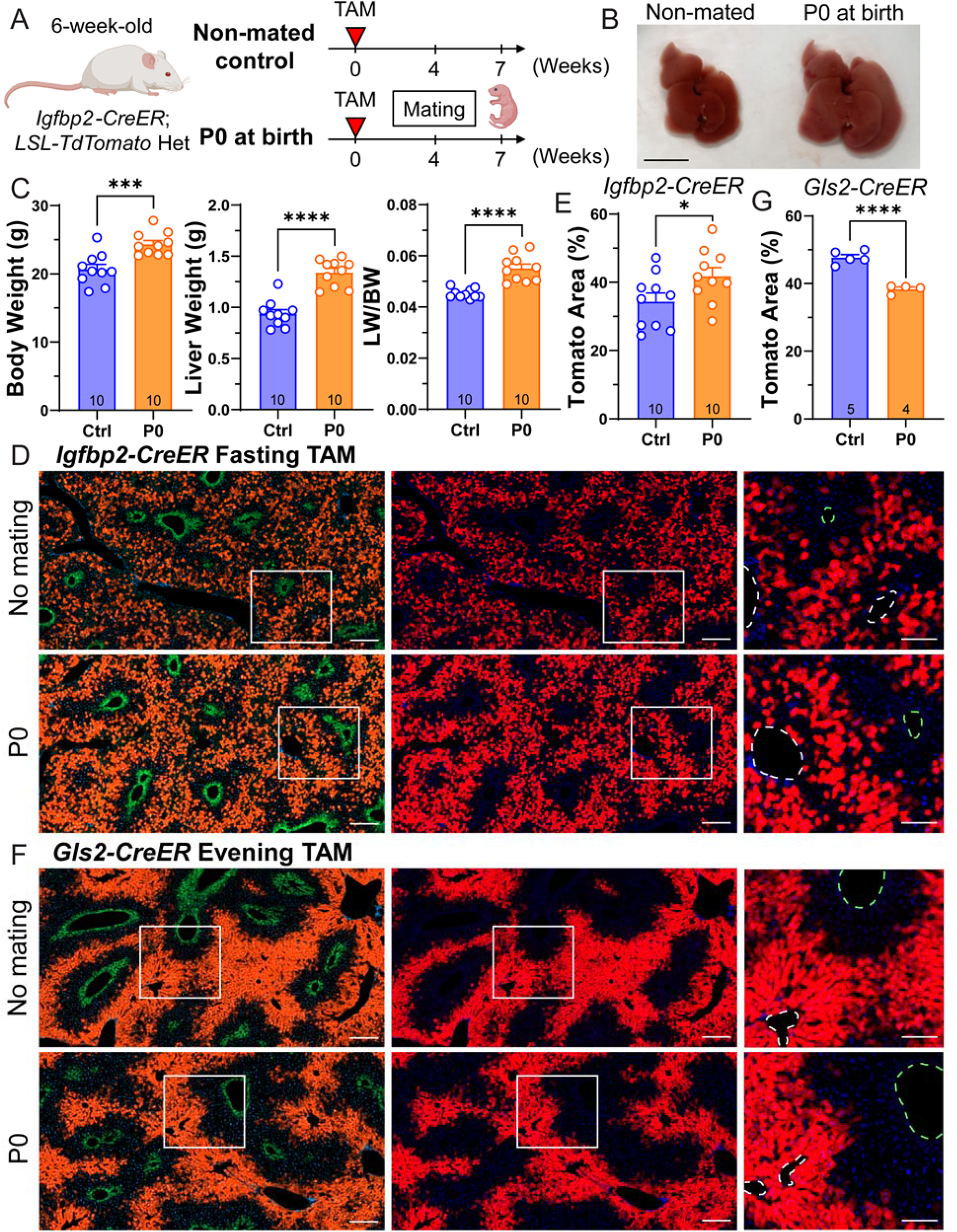
Zone 2 cells contribute to liver expansion during pregnancy. **A**. Schema of lineage tracing experiment during pregnancy. *Igfbp2-CreER* mice were given tamoxifen at 6 weeks of age, and mated 4 weeks after tamoxifen administration. The livers were harvested when the pups were born (P0) in the pregnancy group. **B**. Gross images of livers from control and pregnant mice. Scale bar, 1 cm. **C**. Body weight, liver weight, and liver-to-body weight ratios of no mating controls (Ctrl) and the pregnant group collected at P0 (P0) (n = 10 and 10 for control and pregnant mice). **D**. Representative images of control and pregnant livers from *Igfbp2-CreER* mice. Scale bar, 200 μm for cropped images and 100 μm for magnified images. Slides were stained for GS (green). The green dashed circles represent CVs (marked by GS), and the white dashed circles represent PVs. **E**. Quantification of the Tomato area from **D** (n = 10 and 10 for control and pregnant mice). **F**. Representative images of control and pregnant livers from *Gls2-CreER* mice. Scale bar, 200 μm for cropped images and 100 μm for magnified images. Slides were stained for GS (green). The green dashed circles represent CVs (marked by GS), and the white dashed circles represent PVs. **G**. Quantification of the Tomato area from **F** (n = 5 and 4 for control and pregnant mice).

To lineage trace an orthogonal zone Cre model, we further utilized the *Gls2-CreER* line to trace zone 1 cells. *Gls2-CreER* mice label zone 1 hepatocytes surrounding the PV region (**Figure 5F**). The proportion of Tomato+ cells in *Gls2-CreER* declined during pregnancy (38.37 ± 1.26%) compared to non-mated control females (47.72 ± 1.93%) (**Figure 5G**). Given that there is overlapping Tomato labeling between *Igfbp2-CreER* and *Gls2-CreER* livers, we reason that the zone 2 cells located closer to the central vein (not labeled by *Gls2-CreER*) contributed more toward liver growth during pregnancy.

### Zonation changes during fasting

Surprisingly, the *Igfbp2-CreER* strain labeled a much larger number of midlobular hepatocytes after fasting (**Figure 1**). Because *Igfbp2* might not be the best or only marker for zone 2 cells, we needed to determine if other zone 2 markers are changing with fasting. While changes in the number of *Igbfp2+* cells might suggest isolated changes in *Igfbp2* itself without concomitant changes in other zonated gene expression programs, it is more likely that zonation domains shifted with fasting. If other zone 2 markers change, it would also raise the larger question of how zonation might change between fed and fasted states. While the liver transcriptome is known to massively change during fasting (Sokolović et al., 2008), it is unknown how gene expression changes during fasting on the single cell level. As whole-body metabolism significantly changes during fasting, we speculated that zone specific metabolic functions may also be altered.

To investigate the impact of the fasted state on liver zonation and zone 2 cells, we performed single nuclear RNA sequencing (snRNA-seq) of livers from normal feeding state and fasting for 24 hours. The UMAP plot of all liver cells showed a clear overlap of endothelial, hepatic stellate, and Kupffer cells between fed and fasted conditions (**Figure 6A**). In stark contrast, hepatocytes showed such large transcriptomic differences that fed and fasted hepatocytes were largely non-overlapping (**Figure 6A**). These data showed that fasting drives transcriptomic changes in hepatocytes but not in non-parenchymal cells (NPCs). Given the large differences between feeding and fasting, we analyzed the hepatocyte population separately from NPC populations (see details in Methods). We were able to define three zones within fed and fasted conditions. Strikingly, we observed large increases in the frequency of zone 2 (37.0% to 53.2%) and zone 3 hepatocytes (from 11.1% to 19.7%), and a decrease of zone 1 hepatocytes (42.4% to 17.8%) (**Figure 6B**). To assess individual genes that are representative of zones, we compared the gene expression of zonal markers between fed and fasted states. We found increases in zone 2 (*Hamp, Igfbp2*) and zone 3 (*Oat*), and decreases in zone 1 (*Arg1, Gls2*) genes in fasting livers (**Figure 6C**). These results showed that the frequency of hepatocytes in different zones change dramatically after fasting. Altogether, the results supported that the *Igfbp2* expansion during fasting was not gene-specific, but rather a broader effect on zonation.

**Figure 6.**
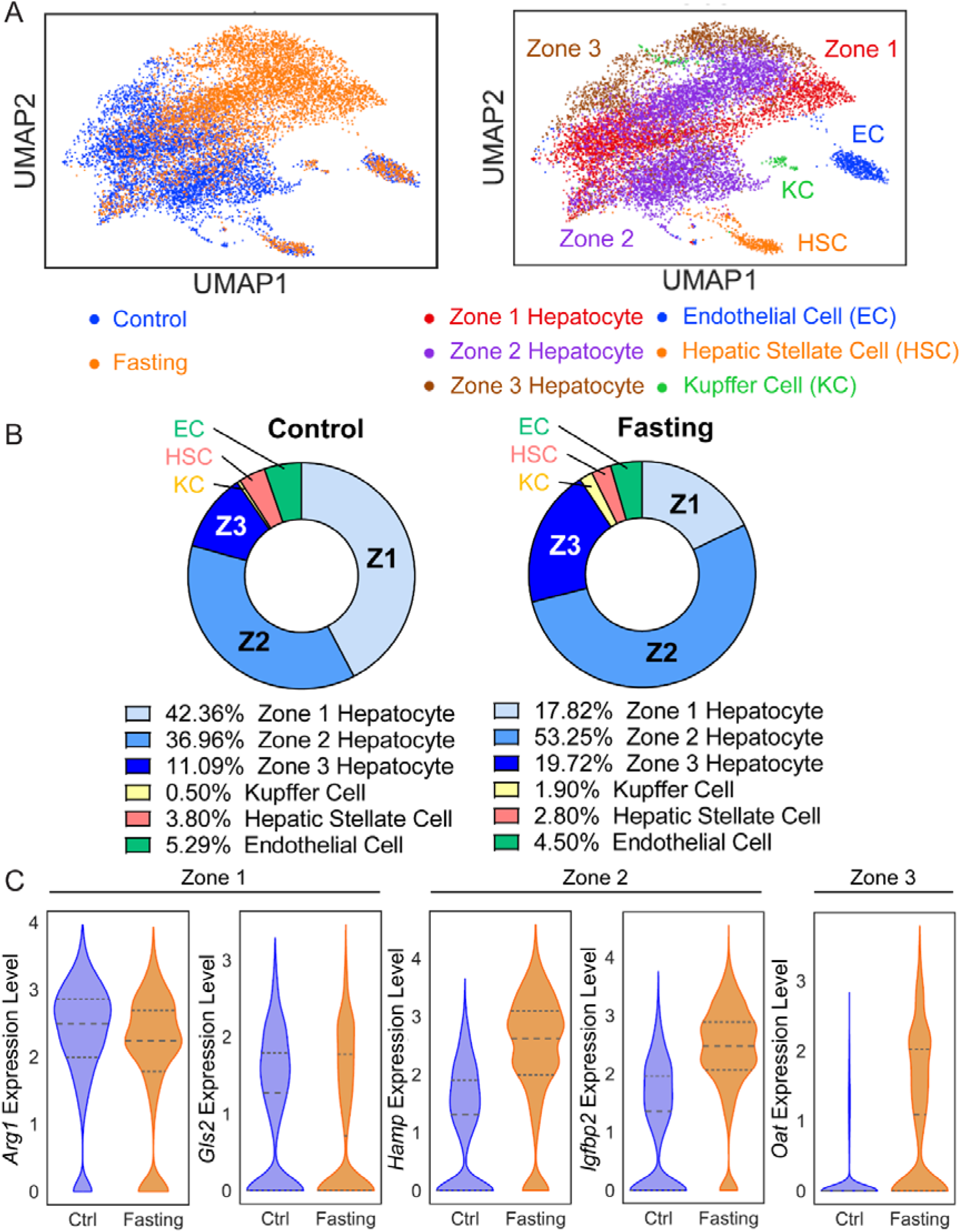
snRNA-seq revealed significant transcriptomic and zonation changes in the fasted liver. **A**. Uniform Manifold Approximation and Projection (UMAP) plots represent the clustering of two samples (left panel) and cell populations (right panel). Nuclei from two fed livers were combined and two fasted livers were combined for the snRNA-seq experiment. **B**. This pie chart indicates the proportions of each cell population in fed (left panel) and fasted livers (right panel). **C**. Gene expression of zonal markers (Zone 1: *Arg1, Gls2*; Zone 2: *Hamp, Igfbp2*; Zone 3: *Oat*) in fed and fasted livers.

Next, we looked for the underlying basis of these zonal changes. Pathways that changed between fed and fasting states were similar in each of the three zones, indicating a coordinated change in metabolic programs throughout the lobule. We observed that oxidative phosphorylation, non-alcoholic fatty liver disease, fatty acid degradation, PPAR signaling pathway, and cholesterol metabolism genes increased with fasting (**Figure S6B**). Overall, all hepatocytes exhibited remarkable transcriptomic changes in the fasted state in order to maintain metabolic homeostasis. Therefore, the transcriptomic differences between zones was diminished. In addition, previously specialized metabolic functions that were more restricted to one zone were no longer restricted, indicating that zonal differences might be suppressed under fasting states. For example, in the fed liver, zone 1 hepatocytes with higher oxygen levels express genes involved in gluconeogenesis (e.g., *Pck1*) and have higher β-oxidation activity (Ben-Moshe and Itzkovitz, 2019). In the fasted liver, all zones were recruited to contribute towards gluconeogenesis and β-oxidation to release stored energy (Rui, 2014). Examination of metabolic pathways in fed or fasting livers showed increased gluconeogenesis and fatty acid oxidation in the fasted state (**Figure S6A**), which could be an explanation for decreased zone 1 population (**Figure 6B**) as other zones are also performing these metabolic functions.

## Discussion

Although previous studies suggested that zone 2 hepatocytes are a major source for liver repopulation (Wei et al. 2021; He et al. 2021; Chen et al. 2020), there has been little direct evidence showing the clonal expansion of this population, and few tools to track or genetically manipulate zone 2 cells specifically. Our results showed that IGFBP2+ cells, which make up about 20-30% of the lobular hepatocyte population, increased in number during homeostasis and expanded more than cells from other zones after many types of injury. After hepatotoxic injuries occurring on either end of the lobule, we observed a regeneration by *Igfbp2*+ cells into the injured area, further confirming that zone 2 hepatocytes are spatially positioned to compensate for cellular loss. Given the controversy in the liver field about the source of new hepatocytes, this additional quantitative evidence for zone 2 hepatocyte regeneration is needed.

There is likely to be additional heterogeneity among hepatocytes in zone 2. First, there is no clear definition of where the boundaries are located between zones. Some studies distinguish zones based on expression of zonal markers (*Cdh* for zone 1, *Gs* for zone 3) (He et al., 2021). We divide the lobule into three compartments based on distance along the PV-to-CV axis. IGFBP2+ cells are mostly in the middle third of the lobule. Based on our zonal definitions, there are some hepatocytes in zone 3 that are GS negative. In this way, we think that not every hepatocyte between the GS+ and the Cdh1+ domains should be considered zone 2.

Even based on our definition of zone 2 being IGFBP2+ cells, there is still heterogeneity within this population. First, the magnitude of expansion of the IGFBP2 population (∼2 fold, from occupying 20.8% to 41.2%) is less than the HAMP2+ population (∼3.7 fold, from 7.4% to 27.4%) during homeostasis over 1 year. Second, lineage tracing during pregnancy showed that *Igfbp2* expressing midlobular cells expanded, while *Gls2* expressing zone 1 cells declined in number. When comparing the occupancy and distribution of Tomato+ cells between these two models, there is considerable overlap in the labeled domains. *Gls2-CreER* labeled approximately half the diameter of the hepatic lobule surrounding the PV, which overlapped with approximately half of the *Igfbp2+* zone 2 cells. This suggested that the zone 2 subpopulation closer to the CV (unlabeled by *Gls2-CreER*) might account for most of the expansion during pregnancy. This is consistent with our previous observation that GLS2+ cells make little contribution to the generation of new hepatocytes under homeostatic conditions (Wei et al., 2021). Nevertheless, both studies came to the same conclusion that midlobular hepatocytes are more proliferative than others in multiple circumstances.

While it is known that heterogeneous metabolic functions are the main feature of hepatic zonation, how metabolic dysregulation changes zonation in physiological or pathological states is unknown. Hepatic gene expression changes dramatically in response to circadian rhythms and fasting/feeding states (Bookout et al. 2013; Sokolović et al. 2008). However, physiological processes that regulate zonation have not been elucidated. In this study, we characterized liver cell populations in normal and fasting states using lineage tracing and snRNA-seq data. The results revealed an increase in zone 2 and zone 3 cells at the expense of zone 1 cells, which correlated with the observation of increased Tomato+ cells in the *Igfbp2-CreER* mice during fasting. It is still an open question whether the zonation change during fasting drives functional outcomes. Interestingly, the pathology of fatty liver disease also progressed in a zonated manner. Are the health benefits of fasting caused in part by increased numbers of hepatocytes taking on zone 2/3 roles? Will the fasted liver become more susceptible to centrilobular damage due to an increase in zone 3 cells? Future work will need to determine how the zonation changes associated with fasting affect liver health and whole body metabolism.

### Limitations of the Study

All the zonation studies were based on mouse models and not human tissues. It is unclear whether hepatic zonation is the same in mouse and human livers. The highly variable labeling efficiency of *Igfbp2-CreER* line may obscure some clonal expansion effects. The transplantation experiments only included two donors, which made statistical analysis difficult. Although the zone 1 and zone 3 compartments are likely changing during fasting, we have not examined the labeling pattern in fasted livers using other zonal *CreER* lines.

## Acknowledgements

We would like to thank D. Ramirez (UTSW Whole Brain Microscopy Facility) for whole liver imaging; J. Xu and Y. J. Kim (CRI Sequencing Core) for sequencing; T.W. is supported by NIH (R01CA258584). H.Z. is supported by the Pollack Foundation, the NIH R01 grants (AA028791, DK125396), a Simmons Comprehensive Cancer Center Cancer & Obesity Translational Pilot Award, and the Emerging Leader Award from the Mark Foundation For Cancer Research (#21-003-ELA).

## Author contributions

Y.H.L. and H.Z. conceived the project, performed the experiments, and wrote the manuscript. Y.W. analyzed the mouse models and performed animal experiments. Y.W., C.A.P. and T.W. performed bioinformatic analysis for the snRNA-seq. M.Z. performed the transplantation surgery. M.Z. and Z.W. generated the scRNA-seq data. M.H.H. assisted with animal experiments and mice husbandry. Y.Z. and T.S. generated the *Igfbp2-CreER* transgenic mouse.

## STAR Methods

### RESOURCE AVAILABILITY

#### Lead Contact and Materials Availability

Further information and requests for resources should be directed to the Lead Contact, Hao Zhu. Mouse lines generated in this study are available from the Lead Contact with a completed Materials Transfer Agreement.

#### Data and code availability

RNA-seq data will be deposited at GEO and made publicly available.

### EXPERIMENTAL MODEL AND SUBJECT DETAILS

#### Animals

The *Igfbp2-CreER* line was generated by CRISPR mouse engineering as described previously (Wei et al., 2021). Briefly, a plasmid containing *IRES-CreERT2* flanked by homology arms from the 3’ UTR of *Igfbp2* was used as a template to generate single stranded (ss)DNAs. SsDNA, CAS9 protein, and sgRNA(s) were co-injected into mouse zygotes. All mice were maintained in a specific pathogen free (SPF) facility and handled in accordance with the guidelines of the Institutional Animal Care and Use Committee (IACUC) of UTSW. Tamoxifen (Sigma-Aldrich #T5648) was dissolved in corn oil and given at a dose of 100 mg/kg into 6-week-old mice as described in **Figure 1**. *FRG* mice were purchased from Yecuris (#10-0001) and maintained on 7.5 μg/mL nitisinone (NTBC, Yecuris #20-0027) water until transplantation.

#### Chemical Injury Experiments

CCl_4_ (Sigma Aldrich #289116) was diluted 1:10 in corn oil and injected one time IP at a dose of 0.5 ml/kg of mouse. Acetaminophen, or APAP (Sigma Aldrich #A7085), was dissolved in warm saline and injected one time IP at 300 mg/kg of mouse. The livers were harvested 2 weeks after injection. DDC was mixed into TestDiet at 0.1% concentration and the mice were fed for 4 weeks starting 2 weeks after tamoxifen.

#### Partial Hepatectomy

Two thirds of the liver was surgically removed in the standard fashion as described previously (Mitchell and Willenbring, 2008). The left lateral lobe and the median lobe were removed, and used as pre-surgical samples for Tomato analysis. The regenerated livers were harvested 2 weeks after surgery. Pre- and post-surgical samples were compared for the same mice using paired Student’s t-tests.

#### Immunofluorescence

Livers were fixed overnight in 4% paraformaldehyde (PFA; Alfa Aesar #J19943K2) at 4°C. For frozen sectioning, the livers were further dehydrated in 30% sucrose overnight at 4°C. Tissues were frozen sectioned at a thickness of 16 μm, washed with PBST three times and blocked in 5% BSA with 0.25% Triton X-100 at room temperature for 1 hour. The slides were incubated with GS antibody (Abcam #ab49873; 1:1000) or FAH antibody (Yecuris #20-0042; 1:500) in the blocking solution at 4°C overnight. After washing three times with PBST, slides were incubated with Alexa Fluor 488 goat anti-rabbit IgG antibody (Life technologies #A-11008) at a 1:200 dilution and Hoechst 33342 (Life technologies #H3570) at 1:1000 dilution in the blocking solution at room temperature for 1 hour. Whole slides were imaged using an Axioscan slide scanner and processed using Zen 2.6 software from Zeiss. ImageJ was used to quantify the percentage of Tomato+ areas as described previously (Wei et al., 2021). Briefly, the composite RBG images were separated into three channels. We then adjusted the threshold to saturate the Tomato signal in the red channel and to cover all the tissue areas in the green channel. Because there is variability across a liver section, Tomato+ areas were calculated using whole slide images rather than individual cropped images. Each mouse in the study had one associated whole slide image.

#### EdU Proliferation Assay

Both the control (*LSL-tdTomato* heterozygous) and *Igfbp2-CreER* (*Igfbp2-CreER; LSL-tdTomato* heterozygous) mice were given tamoxifen at 6 weeks of age. Two weeks after tamoxifen, water containing 1 mg/mL EdU (Carbosynth #NE08701) was provided to males for 14 days. The livers were then harvested and frozen-sectioned. The Click-iT EdU Alexa Fluor 488 Imaging Kit (Life Technologies #C10337) was used to detect the EdU signal. Following EdU staining, GS and Alexa Fluor 647 goat anti-rabbit IgG antibodies (Life technologies #A-21244) were used to mark hepatocytes around CVs. The zonal position indices (P.I.) for EdU+ hepatocytes were determined as previously described (Lin et al., 2018). We only analyzed proliferating hepatocytes, which have large and round nuclei in comparison to the nuclei of non-parenchymal cells. The distance of an EdU+ nuclei to the closest CV (*x*) and closest PV (*y*), and the distance between the CV and PV (*z*) were measured. The position index (P.I.) was calculated using this formula: P.I. = (x^2^ + z^2^ - y^2^)/(2z^2^).

#### Hepatocyte isolation and transplantation

Primary hepatocytes were isolated by the 2-step collagenase perfusion through the inferior vena cava. Mice were perfused with Liver Perfusion Medium (Gibco #17701038) followed by Liver Digest Medium (Gibco #17703034). The liver cell suspension was filtered through a 70 μM strainer. Equal amounts of Hepatocyte Wash Medium (Gibco #17704024) was added to the liver cell suspension and centrifuged at 50 g for 5 minutes to collect hepatocytes. After two washes, the primary hepatocytes were counted and diluted in Hepatocyte Wash Medium. Finally, 1×10^6^ cells were resuspended in 200 μL and transplanted into FRG mice through splenic injection. NTBC water was withdrawn immediately after injection.

#### Single nuclear RNA sequencing (snRNA-seq)

6 week old C57BL/6 mice were purchased from Jackson Laboratory and maintained until 8 weeks of age. Mice were fasted for 24 hours before hepatocyte isolation. Littermates with ad libitum feeding were collected at the same time. After washes, primary hepatocytes were resuspended in GBSS (Sigma Aldrich #G9779) with 0.7% CHAPS (Sigma Aldrich #C5070) and 0.2U/μL RNase inhibitor (Roche Diagnostics #3335399001), and kept on ice for 5 min. The samples were centrifuged at 500g for 5 minutes and further washed twice using PBS with 1% BSA and 0.2U/μL RNase inhibitor. The nuclear extraction efficiency was determined with trypan blue under a microscope. The isolated nuclei were resuspended in the wash buffer to around 1×10^6^ nuclei/mL. Single nuclei libraries were prepared using the 10x Genomics Chromium Single Cell 3’ Reagents Kit v3.1 according to the manufacturer’s protocol. For each sample, two biological replicates (n = 2 mice) were collected and combined. About 8000 nuclei were mixed with reverse transcription master mix and loaded onto Next GEM Chip-G to target ∼5,000 nuclei after recovery. Libraries were sequenced using 150 bp paired-end Illumina NextSeq500 system at the UTSW Children’s Research Institute Sequencing Facility.

#### SnRNA-seq analysis

We used the “mkfastq”, “count” and ‘aggr’ commands to process the 10x scRNA-seq output into one cell by gene expression count matrix, using default parameters. A custom-built version of mouse genome assembly GRCm38 which included unspliced pre-mRNA was used. scRNA-seq data analysis was performed with the Scanpy (1.6.0, ref 2) package in Python (Wolf et al. 2018). Genes expressed in fewer than 3 cells were removed from further analysis. Cells expressing less than 100 and more than 6000 genes were also removed from further analysis. In addition, cells with a high (>= 0.1) mitochondrial genome transcript ratio were removed. For downstream analysis, we used count per million normalization (CPM) to control for library size differences in cells and transformed those into log(CPM+1) values. After normalization, we used the ‘pp.highly_variable_genes’ command in Scanpy to find highly variable genes across all cells using default parameters except for “min_mean = 0.0125 and min_disp=0.5”. The data were then z-score normalized for each gene across all cells. We then used the ‘tl.pca (default parameters)’, the ‘pp.neighbors (n_neighbors=25)’ and the ‘tl.leiden (resolution = 0.2)’ commands in Scanpy to partition the single cells into distinct clusters. Briefly, these processes first identify 50 principle components in data based on the previously found highly variable genes to reduce the dimensions in the original data, and then build a nearest neighbor graph based on the top 25 principle components, and finally a partition of the graph that maximizes modularity was found with the Leiden algorithm (Traag et al. 2019). The most highly expressed genes from each cluster were found through contrasting a cluster of cells with all the other cells (self-vs-rest) using the Wilcoxon signed-rank test, which is included in the “tl.rank_genes_groups” function in the Scanpy package.

##### Statistical analysis

The data in most panels reflect multiple experiments performed on different days using mice derived from different litters. Variation is indicated in the plots using standard error presented as mean ± SEM. In the text, variation is indicated using standard deviation presented as mean ± SD. Two-tailed Student’s t-tests (two-sample equal variance) were used to test the significance of differences between two groups unless otherwise specified. In all figures, statistical significance is displayed as * (p < 0.05), ** (p < 0.01), *** (p < 0.001), ****(p < 0.0001). In the PHx experiment, paired Student’s t-tests were used to test the significance comparing the excised and regenerated liver lobes. The statistical significance is displayed as # (p < 0.05), ## (p < 0.01) in the paired analysis. In all experiments, mice were not excluded from analysis after the experiment was initiated, unless the mice died. Image analysis for quantification of was blinded.

---

**Supplemental Figure S1.**
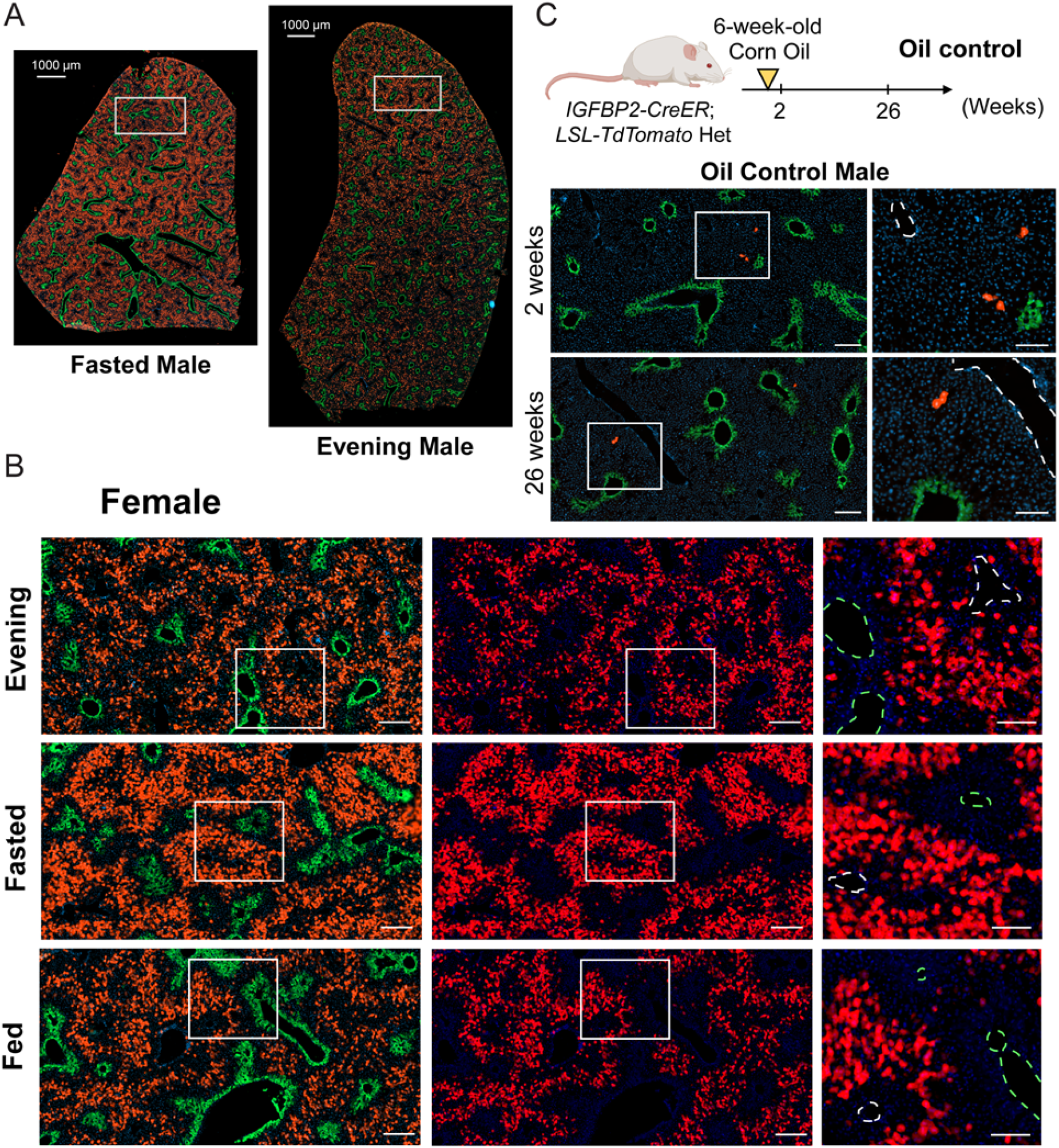
The *Igfbp2-CreER* strain labels midlobular zone 2 hepatocytes after tamoxifen. Data related to Figures 1. **A**. Whole section images of fed tamoxifen and evening tamoxifen males from **Figure 1C**. Scale bar = 1000 μm for cross-sectional images. Slides were stained for GS (green). **B**. Representative images of female *Igfbp2-CreER* labeling by three distinct tamoxifen approaches. Scale bar = 200 μm for cropped images and 100 μm for magnified images. Slides were stained for GS (green). The green dashed circles represent CVs (marked by GS), and the white dashed circles represent PVs. **C**. Schema and representative images of *Igfbp2-CreER; LSL-tdTomato* livers 2 and 26 weeks after oil administration without tamoxifen. Scale bar = 200 μm for cropped images and 100 μm for magnified images. Slides were stained for GS (green). The white dashed circles represent PVs.

**Supplemental Figure S2.**
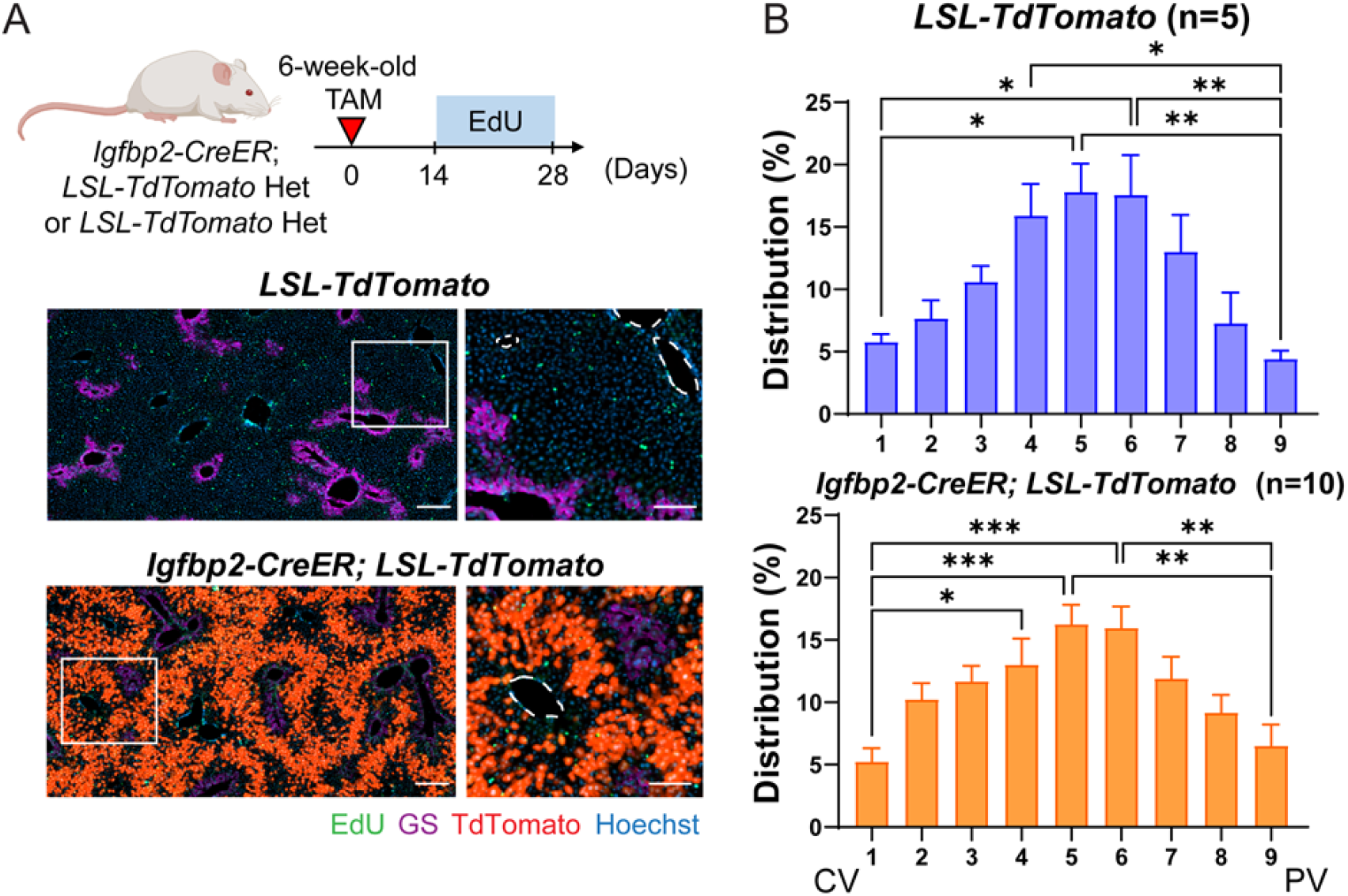
The *Igfbp2-CreER* transgene does not alter the distribution of zonal proliferation when compared to WT littermates. Data related to Figures 1. **A**. Schema and representative images of *LSL-tdTomato* and *Igfbp2-CreER; LSL-tdTomato* after fasting tamoxifen followed by 14 days of EdU incorporation. Scale bar = 200 μm for cropped images and 100 μm for magnified images. Slides were stained for EdU (green) and GS (purple). The white dashed circles represent PVs. **B**. Quantification of spatial distribution of EdU+ cells from **A**.

**Supplemental Figure S3.**
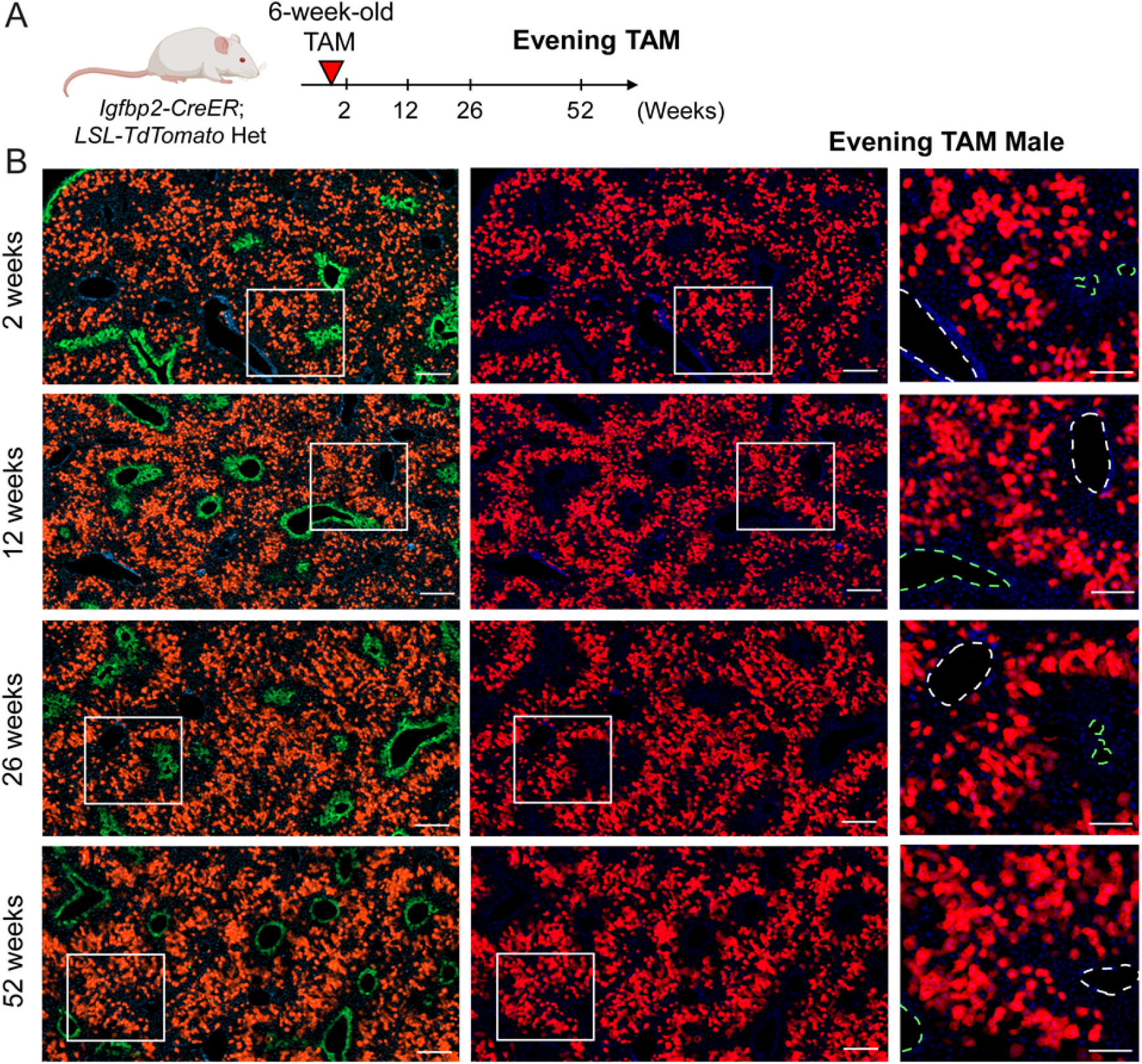
Zone 2 cells labeled by the evening tamoxifen approach expand during homeostasis in female mice. Data related to Figure 2. **A**. Schema of evening tamoxifen lineage tracing experiment under normal homeostasis. **B**. Representative images of male evening tamoxifen tracing over 2, 12, 26 and 52 weeks. Scale bar, 200 μm for cropped images and 100 μm for magnified images. Slides were stained for GS (green). The green dashed circles represent CVs (marked by GS), and the white dashed circles represent PVs.

**Supplemental Figure S4.**
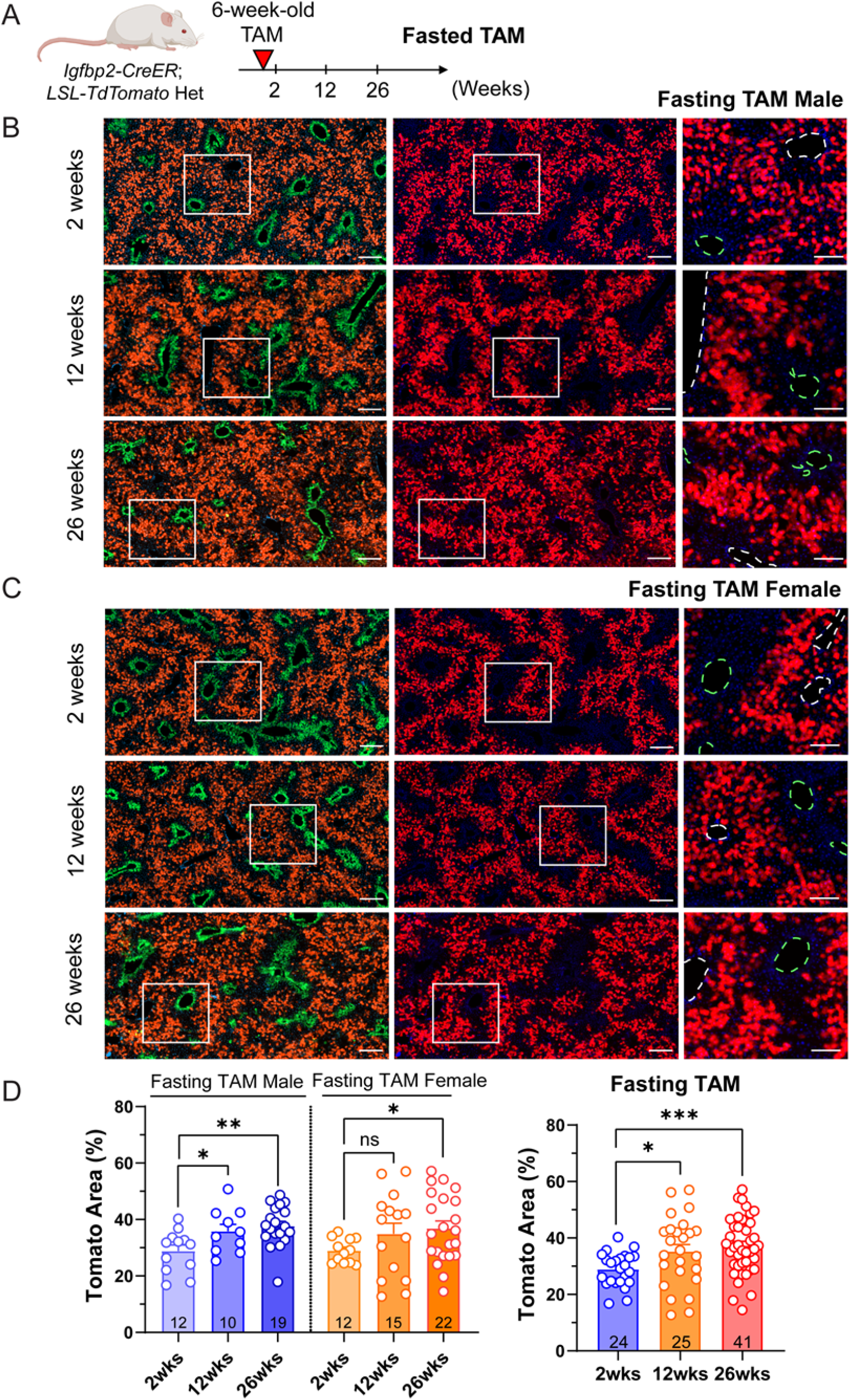
Zone 2 cells expand during homeostasis. Data related to Figures 2. **A**. Schema of fasting tamoxifen lineage tracing experiment under normal homeostasis. **B**. Representative images of male fasting tamoxifen tracing over 2, 12 and 26 weeks. Scale bar, 200 μm for cropped images and 100 μm for magnified images. Slides were stained for GS (green). The green dashed circles represent CVs (marked by GS), and the white dashed circles represent PVs. **C**. Representative images of female fasting tamoxifen tracing over 2, 12 and 26 weeks. Scale bar, 200 μm for cropped images and 100 μm for magnified images. Slides were stained for GS (green). The green dashed circles represent CVs (marked by GS), and the white dashed circles represent PVs. **D**. Quantification of the Tomato area from **B** and **C**. The right panel combines all of the data from the two sexes shown in the left panel. The 2 week data points are the same data as the fasting timepoints in **Figure 1E** (n = 24, 25, 41 mice for 2, 12 and 26 weeks).

**Supplemental Figure S5.**
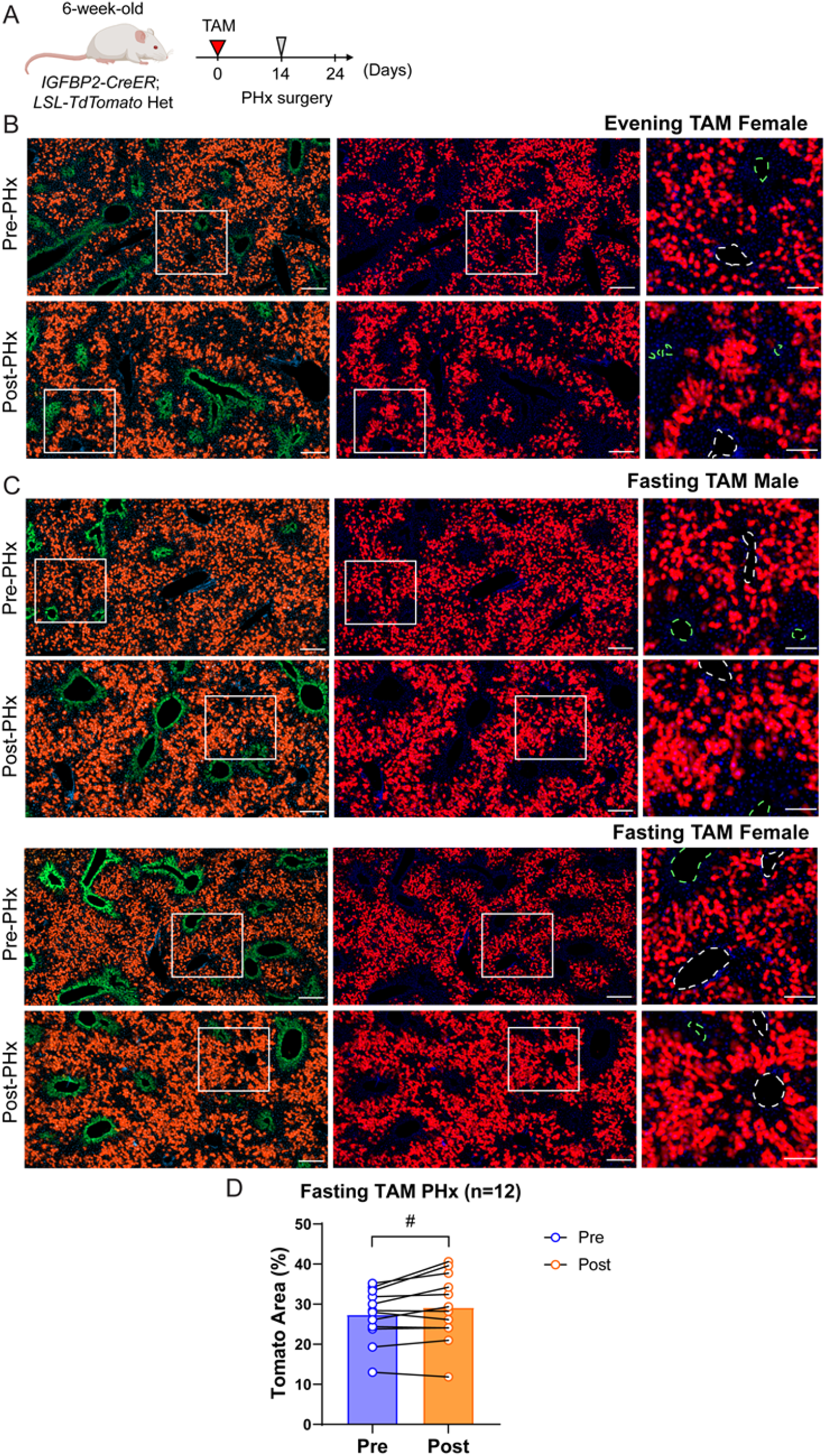
Zone 2 hepatocytes labeled in *Igfbp2-CreER* mice expand after PHx. Data related to Figure 4. **A**. Schema of the *Igfbp2-CreER* lineage tracing experiment in the context of PHx. **B**. Representative images of evening tamoxifen labeled females, pre- and post-PHx. Scale bar, 200 μm for cropped images and 100 μm for magnified images. Slides were stained for GS (green). The green dashed circles represent CVs (marked by GS), and the white dashed circles represent PVs. **C**. Representative images of fasting tamoxifen pre- and post-PHx. Scale bar, 200 μm for cropped images and 100 μm for magnified images. Slides were stained for GS (green). The green dashed circles represent CVs (marked by GS), and the white dashed circles represent PVs. **D**. Quantification of the Tomato area from **C**. Significance was assessed by paired t-tests (n = 12 mice).

**Supplemental Figure S6.**
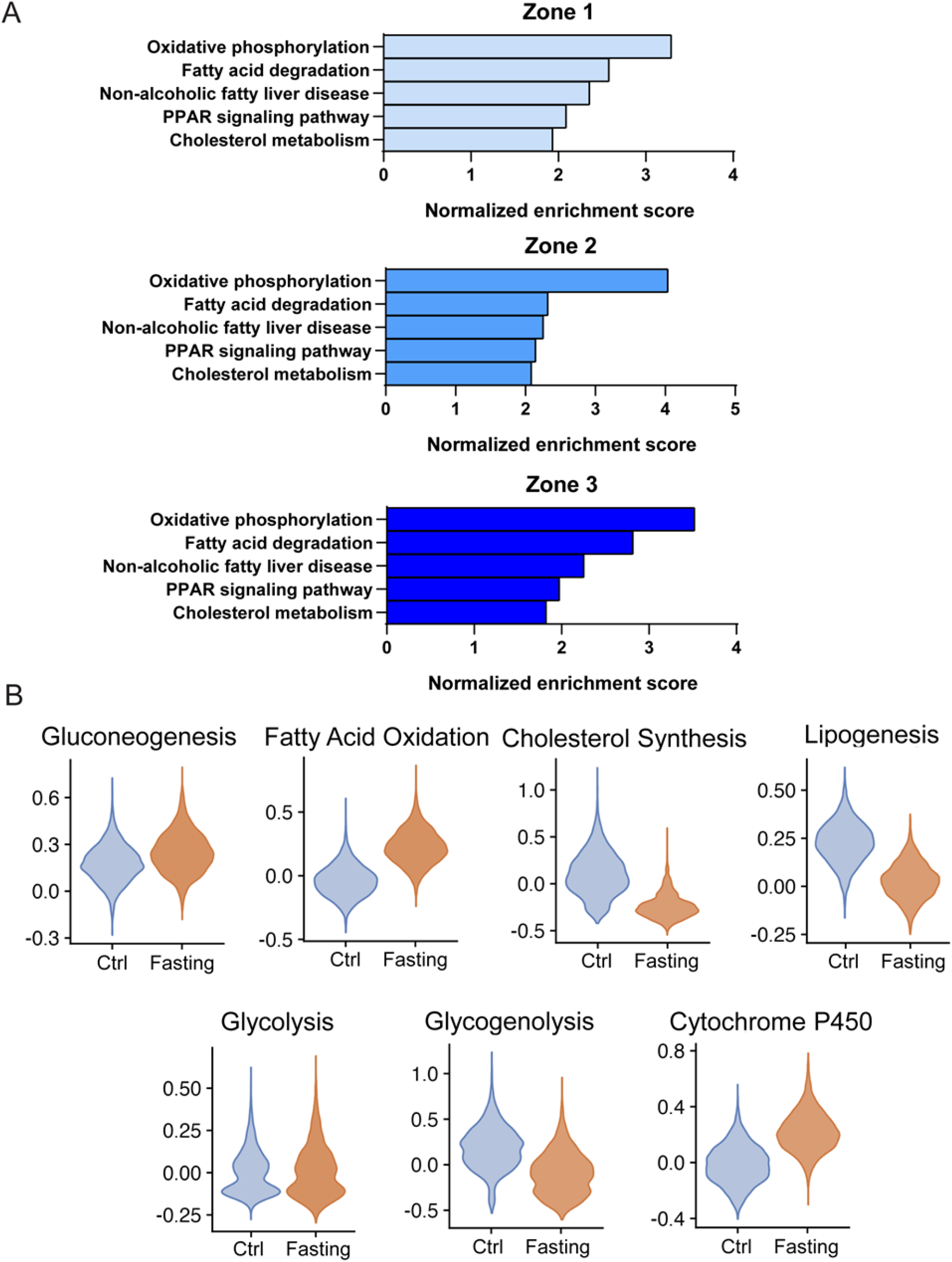
Metabolic pathways change in the fasted liver. Data related to Figure 6. **A**. KEGG pathway enrichment analysis of differentially expressed genes in each zonal population. **B**. Metabolic pathway analysis of the snRNA-seq dataset grouped by normal and fasting conditions.

